# Enhancement of nitrous oxide emissions in soil microbial consortia via copper competition between proteobacterial methanotrophs and denitrifiers

**DOI:** 10.1101/2020.09.21.307504

**Authors:** Jin Chang, Daehyun Daniel Kim, Jeremy D. Semrau, Juyong Lee, Hokwan Heo, Wenyu Gu, Sukhwan Yoon

## Abstract

Unique means of copper scavenging have been identified in proteobacterial methanotrophs, particularly the use of methanobactin, a novel ribosomally synthesized post-translationally modified polypeptide that binds copper with very high affinity. The possibility that copper sequestration strategies of methanotrophs may interfere with copper uptake of denitrifiers *in situ* and thereby enhance N_2_O emissions was examined using a suite of laboratory experiments performed with rice paddy microbial consortia. Addition of purified methanobactin from *Methylosinus trichosporium* OB3b to denitrifying rice paddy soil microbial consortia resulted in substantially increased N_2_O production, with more pronounced responses observed for soils with lower copper content. The N_2_O emission-enhancing effect of the soil’s native *mbnA*-expressing *Methylocystaceae* methanotrophs on the native denitrifiers was then experimentally verified with a *Methylocystaceae*-dominant chemostat culture prepared from a rice paddy microbial consortium as the inoculum. Lastly, with microcosms amended with varying cell numbers of methanobactin-producing *Methylosinus trichosporium* OB3b before CH_4_ enrichment, microbiomes with different ratios of methanobactin-producing *Methylocystaceae* to gammaproteobacterial methanotrophs incapable of methanobactin production were simulated. Significant enhancement of N_2_O production from denitrification was evident in both *Methylocystaceae*-dominant and *Methylococcaceae*-dominant enrichments, albeit to a greater extent in the former, signifying the comparative potency of methanobactin-mediated copper sequestration while implying the presence of alternative copper abstraction mechanisms for *Methylococcaceae*. These observations support that copper-mediated methanotrophic enhancement of N_2_O production from denitrification is plausible where methanotrophs and denitrifiers cohabit.

**Importance:** Proteobacterial methanotrophs, groups of microorganisms that utilize methane as source of energy and carbon, have been known to utilize unique mechanisms to scavenge copper, namely utilization of methanobactin, a polypeptide that binds copper with high affinity and specificity. Previously the possibility that copper sequestration by methanotrophs may lead to alteration of cuproenzyme-mediated reactions in denitrifiers and consequently increase emission of potent greenhouse gas N_2_O has been suggested in axenic and co-culture experiments. Here, a suite of experiments with rice paddy soil slurry cultures with complex microbial compositions were performed to corroborate that such copper-mediated interplay may actually take place in environments co-habited by diverse methanotrophs and denitrifiers. As spatial and temporal heterogeneity allow for spatial coexistence of methanotrophy (aerobic) and denitrification (anaerobic) in soils, the results from this study suggest that this previously unidentified mechanism of N_2_O production may account for significant proportion of N_2_O efflux from agricultural soils.

## Introduction

Methane (CH_4_) and nitrous oxide (N_2_O) are the most influential greenhouse gases apart from CO_2_, with estimated contributions of 16% and 6.2% to global greenhouse gas emissions over a 100-year-time frame (1). Biological sources and sinks significantly impact both atmospheric CH_4_ and N_2_O pools. That is, methanotrophs consume much (up to 100%) of CH_4_ originating from methanogenesis, balancing the global CH_4_ budget (2). Similarly, N_2_O produced via microbially-mediated nitrification and denitrification is offset by N_2_O reduction mediated by microbes capable of expressing and utilizing nitrous oxide reductases (NosZ) (3-5). Collectively, the relative abundance and activity of different microbial groups are critical in determining if any particular environment is a net source or sink of these potent greenhouse gases.

Methanotrophy has become a rather comprehensive term following recent discovery of extremophilic verrucomicrobial methanotrophs, nitrite-reducing anaerobic NC-10 methanotrophs, and archaeal anaerobic methanotrophs (6-9); however, except for certain specialized extreme habitats, “conventional” aerobic proteobacterial methanotrophs often dominate methane oxidation *in situ*, especially in terrestrial systems, e.g., oxic-anoxic interfaces of rice paddy soils and landfill cover soils, as observed in recent metagenomic analyses (10, 11). Proteobacterial methanotrophs are phylogenetically subdivided into gammaproteobacterial and alphaproteobacterial subgroups (12). Both of these organismal groups utilize particulate and/or soluble methane monooxygenases (MMO) encoded by *pmo* and *mmo* operons, respectively, and these MMOs are currently known as the only enzymes capable of preferentially catalyzing the oxidation of CH_4_ to CH_3_OH (13). Of the two forms of MMOs, the particulate methane monooxygenase (pMMO) has been regarded as the prevalent form in most terrestrial environments, and the majority of proteobacterial methanotroph genomes sequenced to date contain only *pmo* operon(s) (14). The expression and activity of pMMO are strongly dependent on copper, although the exact role of copper in pMMO still remains unanswered (12, 15, 16).

Due to the dependency of pMMO activity on copper, pMMO-expressing methanotrophs have high demands for copper (17). Not surprisingly, methanotrophs have developed effective copper uptake mechanisms, presumably to cope with limited copper availability *in situ*. Some alphaproteobacterial methanotrophs of the *Methylocystaceae* family produce and excrete methanobactin, a modified peptide ∼800-1300 Da in size that chelates copper with the highest affinity among known metal chelators (empirical copper-binding constants range between 10^18^-10^58^ M^-1^ depending on experimental protocol used) (18). Methanobactin has high specificity towards copper, and binding constants higher than 10^9^ M^-1^ have been observed only for its complexation with Cu^+^, Cu^2+^ (reduced to Cu^+^ upon binding), Au^3+^, and Hg^2+^ (17). Copper-bound methanobactin is transported into the cell via a TonB-dependent transporter and presumably utilized for synthesis of a functional pMMO complex (19, 20). The genes encoding for the methanobactin polypeptide precursor (*mbnA*) and the enzymes involved in its post-translational modifications have been identified in many alphaproteobacterial methanotroph genomes (roughly half of >40 genomes currently available in the NCBI database), and methanobactin isolated from seven distinct strains of *Methylocystaceae* methanotrophs have been chemically characterized, suggesting that the capability to synthesize and utilize methanobactin is a wide-spread, but not universal, trait for *Methylocystaceae* methanotrophs (18, 21-24).

In CH_4_-rich, copper-depleted environments, these copper acquisition mechanisms of methanotrophs may interfere with other biogeochemical reactions catalyzed by cuproenzymes. Several key enzymes involved in the biological nitrogen cycle including bacterial and archaeal ammonia monooxygenases (AMO), copper-dependent NO_2_^-^ reductases (NirK), and N_2_O reductases (NosZ) require copper ions for their activity (25-27). The impact on NosZ expression and activity would be particularly consequential from an environmental perspective, as NosZ-catalyzed N_2_O reduction is the only identified biological or chemical N_2_O sink in the environment (3, 28). Ammonia oxidation mediated by bacterial AMO retains its activity at copper-deficient incubation condition (i.e., nanomolar bioavailable Cu concentration), and NO_2_^-^ reduction to NO in denitrifiers can be carried out by Cu-independent cytochrome *cd*_1_ nitrite reductase (NirS); however, no copper-independent alternative pathway for N_2_O reduction to N_2_ has been identified to date (3, 29). In fact, methanobactin-mediated inhibition of NosZ activity was recently experimentally verified *in vitro* using simple, well-defined co-cultures of *M. trichosporium* OB3b and several denitrifier strains possessing *nosZ*, suggesting the possibility of increased N_2_O emissions *in situ* where methanotrophs and denitrifiers co-exist (30). Oxic-anoxic interfaces and the vadose zones with fluctuating water content would provide settings in organic- and nitrogen-rich soils, where co-occurrence of obligately aerobic methanotrophy and obligately anaerobic denitrification is possible (31, 32). Whether such an N_2_O-production mechanism is truly relevant in the field, however is not yet known, as neither methanobactin production nor its influence on denitrifiers has been observed in complex microbiomes such as agriculture soils.

As a follow-up to our previous study documenting the impact of methanobactin-producing methanotroph on denitrification and N_2_O production in simple co-cultures, the current study investigated further the potential ecological relevance of this methanotroph-denitrifier interaction by introducing microbial complexity and competition into the picture. The susceptibility of soil’s complex denitrifying consortia to methanobactin-mediated alteration was examined with (1) denitrifying enrichments amended with exogenous addition of methanobactin and (2) stimulation of native alphaproteobacterial methanotrophs of the *Methylocystaceae* family. Further, the consequence of alphaproteobacterial methanotrophs-vs-gammaproteobacterial methanotrophs competition on denitrification and associated N_2_O production was examined with soil slurry microcosms augmented with varying amounts of *M. trichosporium* OB3b before CH_4_ enrichment.

## Results

### The effect of methanobactin from *M. trichosporium* OB3b on N_2_O production in denitrifying soil enrichments

The effect of the methanobactin isolated from *M. trichosporium* OB3b (MB-OB3b) on N_2_O production was examined with NO_3_^-^-reducing enrichments of five rice paddy soils with varying copper content (Fig. 1; Supplementary Table S1). Without added MB-OB3b, the amount of accumulated N_2_O-N did not exceed 0.2% of ∼250 µmoles NO_3_^-^ initially added to the reaction vessels at any time in any of the enrichment cultures despite four out of five soil samples being slightly acidic with 5.8<pH<6.3 (Fig. 1A-E). In contrast, substantial N_2_O accumulation was observed in soil slurry cultures amended with 10 µM MB-OB3b, with the exception of soil with the highest copper content (Fig. 1F-J). Permanent N_2_O accumulation was observed in the enrichments prepared with the soils with the lowest copper content (A: 1.14±0.06 mg Cu/kg dry wt; B: 1.50±0.02 mg Cu/kg dry wt), with 138±2 and 102±11 µmoles N_2_O-N produced from denitrification of 257±29 and 275±5 µmoles NO_3_^-^, respectively. Although the non-stoichiometric N_2_O production suggested partial N_2_O reduction activity, no further N_2_O reduction was observed for at least 24 h after completion of denitrification in either enrichment. Inhibition of NO_2_^-^ reduction was also evident in the Soil B enrichment (Fig. 1G). Permanent N_2_O accumulation and delayed NO_2_^-^ reduction was observed also with the enrichment prepared with Soil C that had a higher copper content (2.78±0.28 mg Cu/kg dry wt; Fig. 1H). The enrichments prepared with soils with the highest copper contents (D: 5.73±0.38 mg Cu/kg dry wt; E: 8.05±0.34 mg Cu/kg dry wt) did not permanently accumulate N_2_O even with 10 µM methanobactin added; however, the maximum amount of transiently accumulated N_2_O was significantly higher (*p*<0.05) with methanobactin than without. No significant NO_3_^-^ or NO_2_^-^ reduction was observed in any of the sterilized controls, and N_2_O production over 120 hours of abiotic incubation yielded <1.0 µmoles N_2_O-N, precluding the possibility that abiotic N_2_O production contributed significantly to N_2_O produced in the NO_3_^-^-reducing soil enrichments (data not shown). These results demonstrated that methanobactin can inhibit N_2_O and/or NO_2_^-^ reduction in complex microbial consortia, but the effect may vary depending on soil properties.

**FIG 1.**
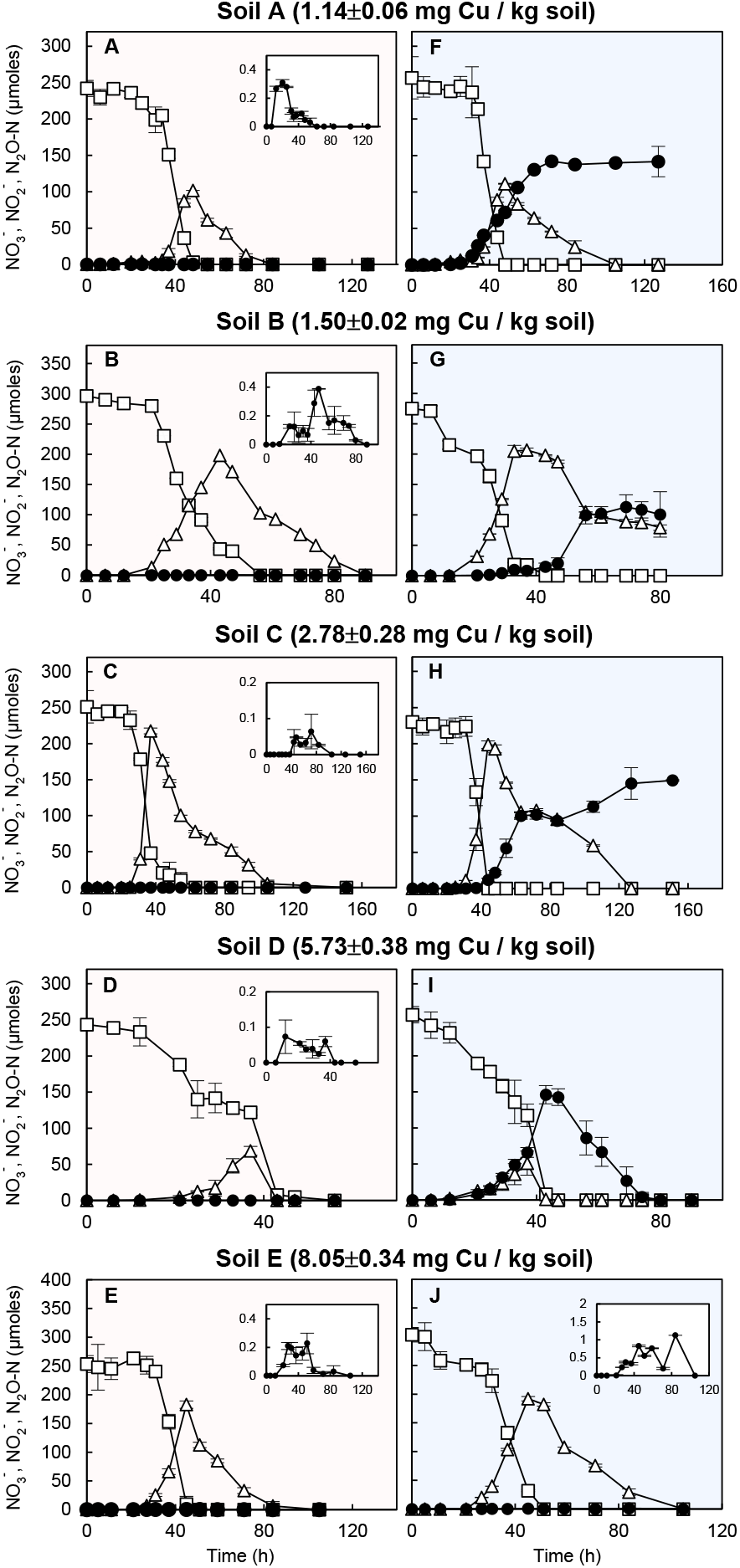
Denitrification of 250 μmoles (5 mM) NO_3_^−^ added to 50 mL rice paddy soil suspensions in 160-mL serum bottles amended without (A-E; shaded red) and with (F-J; shaded blue) 10 μM OB3b-MB. The copper content of the rice paddy soils used for preparation of the suspensions were 1.14 (A, F), 1.50 (B, G), 2.78 (C, H), 5.73 (D, I), and 8.05 (E, J) mg Cu / kg dry wt. The time series of the average amounts (μmoles per vessel) of NO_3_^−^ (□), NO_2_^−^ (△), and N_2_O (●) are presented with the error bars representing the standard deviations of triplicate samples. The inserts are magnification of the N_2_O monitoring data.

### Denitrification and N_2_O accumulation in a soil microbial consortium enriched with indigenous *Methylocystaceae* methanotrophs

To observe whether indigenous soil methanotrophs are capable of altering soil denitrification and enhancing N_2_O production with methanobactin as the mediator, denitrification was observed with *Methylocystaceae*-dominant quasi-steady state culture extracted from a chemostat (Fig. 2, Supplementary Table S2). The copy number of eubacterial 16S rRNA genes in the quasi-steady state reactor culture was 1.52±0.02×10^6^ copies mL^-1^. Alphaproteobacterial methanotrophs of the *Methylocystaceae* family were the dominant bacterial population, with 78 % of the 16S rRNA reads assigned to OTUs affiliated with this taxon, while the OTUs assigned to the gammaproteobacterial methanotrophs (the *Methylococcaceae* family) constituted a minority group with 1.5 % relative abundance. Non-methanotrophic taxa identified in the reactor culture included *Chitinophagaceae* (7.3%), *Pseudomonadaceae* (1.4%), and *Mycobacteriaceae* (1.3%). The most abundant *nirK* and *nosZ* recovered from the metagenome of the chemostat culture were most closely affiliated in terms of translated amino acid sequences to *Methylocystis* sp. Rockwell (69 % of recovered *nirK*) and *Methylocystis* sp. SC2 (93 % of recovered *nosZ*), both of which are the only *nirK* and *nosZ* found in sequenced alphaproteobacterial methanotroph genomes (Supplementary Table S3). The other *nirK* genes recovered with >1% relative abundance (among the recovered *nirK* sequences) included those most closely affiliated to the genera *Mesorhizobium* (16 %), *Bauldia* (3.5%), *Panacibacter* (3.1%), *Pseudomonas* (1.8%), and *Hypomicrobium* (1.4%). The most abundant non-methanotrophic *nosZ* genes included those affiliated to the genera *Flavobacterium* (3.4%; clade II) and *Pseudomonadas* (2.5%; clade I). The coverage of *nirS* genes were substantially lower than those of *nirK* (2.1 × 10^−3^ *nirS/recA* as compared to 0.133 total *nirK*/*recA* and 4.2 × 10^−2^ non-*Methylocystis nirK*/*recA*). The recovered *nirS* genes included those affiliated to the genera *Bradyrhizobium* (29.0% of recovered *nirS*), *Zoogloea* (41.0%), and *Acidovorax* (31.9%).

**FIG 2.**
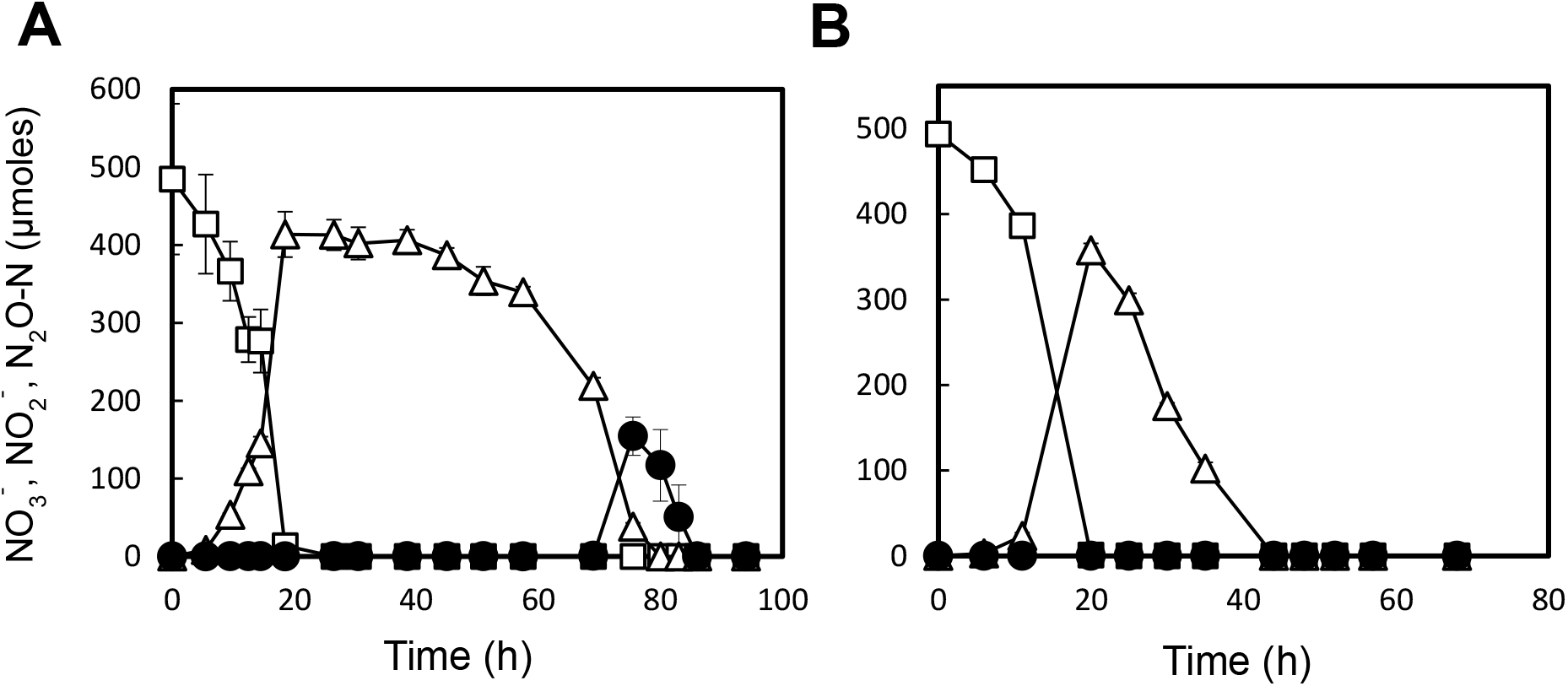
(A) Progression of denitrification in the 50-mL enrichment cultures (in 160-mL serum bottles with N_2_ headspace) extracted from the *Methylocystaceae*-dominant chemostat. One milliliter of separately prepared denitrifier enrichment culture was added to the enrichment cultures at t=0. To a set of controls (B), CuCl_2_ was added to a concentration of 2 µM before incubation. The changes to the amounts of NO_3_^−^ (□), NO_2_^−^ (△), and N_2_O (●) in the culture vessels were monitored. The error bars represent the standard deviations of triplicate samples.

The sole unique *mbnA* sequence assembled from the shotgun metagenome reads exhibited high similarity to *mbnA* sequences of organisms affiliated to the *Methylocystis* genus (Supplementary Fig. S1). The qPCR quantification targeting the *Methylocystaceae mbnA* genes estimated 8.1±1.8×10^4^ copies mL^-1^, which translated to an *mbnA*-to-16S rRNA ratio of 0.054. Further, the *mbnA* transcript-to-gene ratio was 19.5±4.9 as determined using RT-qPCR, indicating that the *mbnA* gene was actively transcribed during quasi-steady state operation of the reactor.

Reduction of NO_3_^-^ was observed immediately after addition of the denitrifying inoculum to the degassed CH_4_-enriched culture extracted from the chemostat and was unaffected by Cu^2+^ amendment (Fig. 2B). Repression of NO_2_^-^ reduction was apparent in the culture without Cu^2+^ amendment, as no significant decrease in NO_2_^-^ concentration was observed for ∼20 hours after maximum NO_2_^-^ accumulation was observed (414±29 µmoles NO_2_ at t=18.5 h). Eventual reduction of NO_2_^-^ led to transient accumulation of N_2_O, and the maximum amount of N_2_O observed before it was presumably reduced to N_2_ was 155±25 µmoles N_2_O-N (∼32% of added NO_3_^-^-N). Such apparent partial repression of NO_2_^-^ and N_2_O reduction was absent in the samples amended with 2 μM Cu^2+^. NO_2_^-^ reduction progressed without any apparent lag, and the amount of N_2_O in the vessel did not increase higher than 0.09±0.01 µmoles N_2_O-N before it was consumed. These observations suggest that alteration of copper availability by methanobactin-producing *Methylocystaceae* methanotrophs had substantial effect on denitrification and N_2_O reduction in the broader microbial community.

### Effects of altered community compositions of methanotrophic enrichments on denitrification and N_2_O production upon transition to anoxia

Methanotroph-enriched microbiomes with different ratios of methanobactin-producing *Methylocystaceae* to gammaproteobacterial methanotrophs were mimicked in soil slurry microcosms by adding varying amounts of *M. trichosporium* OB3b precultures to rice paddy soil slurries before batch enrichment with 20% v/v CH_4_ (Fig. 3). The 16S rRNA gene copy number of *Methylococcaceae* methanotrophs in the soil sample was estimated to be 6.5±1.2×10^6^ copies g wet wt soil^-1^ from their relative abundance (0.41%) and the total bacterial 16S rRNA copy number (1.6±0.3×10^9^ bacterial 16S rRNA copies g wet wt soil^-1^). *Methylosinus trichosporium* OB3b *pmoA* copy numbers in the CH_4_-enriched slurries were 0, 1.2±0.2×10^6^, and 3.3±0.7×10^8^ per mL in the cultures that had been augmented with *M. trichosporium* OB3b *pmoA* copy numbers in quantities matching 0, 1, and 10 times the estimated 16S rRNA gene copy numbers affiliated to *Methylococcaceae* methanotrophs (referred to as *mtri0, mtri1*, and *mtri10*, respectively). The 16S rRNA amplicon sequencing of the same samples estimated the relative abundances of *Methylocystaceae* methanotrophs to be 0.15, 0.35 and 13.0%, respectively (Supplementary Table S4). The relative abundances of *Methylococcaceae* methanotrophs were 39, 7.7, and 0%, respectively, in the same samples.

**FIG 3.**
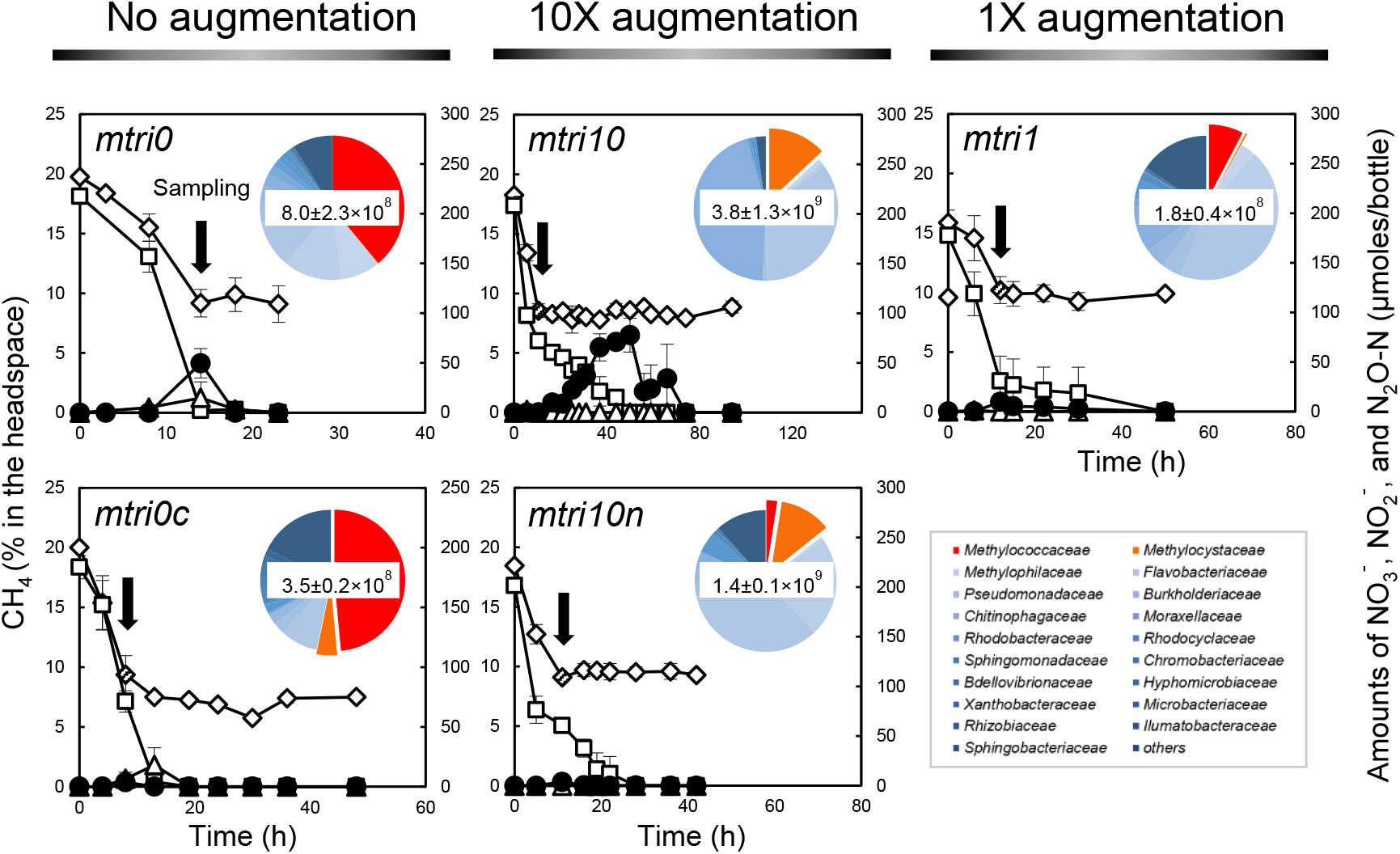
Monitoring of CH_4_ oxidation and subsequent denitrification and N_2_O production in rice paddy soil suspensions augmented with *M. trichosporium* OB3b (or Δ*mbnAN* mutant) cells prior to incubation to quantities with their *pmoA* copy numbers matching 0 (*mtri0, mtri0c*), 1 (*mtri1*), and 10 *(mtri10, mtri10n*) times the estimated *Methylococcaceae* 16S rRNA gene copy numbers in the suspensions. The Cu-replete control, to which 2 µM CuCl_2_ was added (*mtri1c*), and the control augmented with Δ*mbnAN* mutant of *M. trichosporium* (*mtri10n*) were included. The time series of the amounts (μmoles per vessel) of NO_3_^−^ (□), NO_2_^−^ (△), and N_2_O (●) and the headspace concentrations (%) of CH_4_ (⋄) are presented with the error bars representing the standard deviations of triplicate samples. The pie charts embedded in the panels illustrate the microbial community compositions of the enrichment cultures at the time points indicated by the black arrows. The numbers in the center of the pie charts are the estimates of bacterial 16S rRNA copy numbers per mL of culture at the indicated time points. The detailed microbial community compositions are presented in Supplementary Table S4.

In the *mtri0* culture, the maximum amount of accumulated N_2_O was 49.6±14.5 µmoles N_2_O-N and the duration of N_2_O accumulation was shorter than 10 h (Fig. 3). The maximum observed N_2_O accumulation (78.2±16.4 µmoles N_2_O-N at t=50 h) was significantly higher (*p*<0.05) and the duration of N_2_O accumulation substantially longer (68.5 h) in the *mtri10* cultures, in which *Methylocystaceae* methanotrophs outnumbered *Methylococcaceae* methanotrophs. The decrease in the amount of NO_3_^-^ observed during active CH_4_ oxidation (t<10.5 h) was presumably due to assimilation, as denitrification was unlikely to occur before O_2_ depletion. Thus, N_2_O accumulation resulting from dissimilatory reduction of NO_3_^-^ between 10.5 and 94 h was near stoichiometric, albeit transient, in the *mtri10* cultures. In the *mtri1* cultures, the maximum amounts of accumulated N_2_O were significantly lower (*p*<0.05) than either the *mtri0* and *mtri10* cultures, in line with the substantially lower methanotroph population.

In the negative control experiment performed with the Δ*mbnAN* mutant strain added to the soil suspension in place of the wildtype *M. trichosporium* OB3b (referred to as *mtri10n*), the maximum amount of N_2_O accumulated in the vessel was limited to 0.27±0.09 µmoles N_2_O-N despite the *Methylocystaceae* dominance of the methanotrophic population, as indicated by the high *M. trichosporium* OB3b *pmoA* copy numbers (1.2±0.7×10^8^ per mL) and the low *Methylococcaceae*-to-*Methylocystaceae* population ratio (0.21) in the *mtri10n* culture (Fig. 3). The results of this control experiment confirmed that the enhanced N_2_O accumulation of the *Methylocysteceae*-dominant enrichment was due to copper sequestration by methanobactin produced by *M. trichosporium* OB3b. Also notable was the obvious dissimilarity observed in the microbial community composition between the enrichments with the wildtype strain OB3b and Δ*mbnAN* mutant strain (Fig. 3, Supplementary Table S4). The additional set of control experiment performed with 2.0 μM CuCl_2_ added to the soil-only, thus *Methylococcaceae*-dominant, samples (referred to as *mtri0c*), suggested that the instantaneous N_2_O accumulation observed in the *Methylococcaceae*-dominant culture could also be explained as the effect of Cu^2+^ sequestration by the methanotrophic population. The maximum N_2_O production (3.75±1.25 µmoles N_2_O-N at t=106 h) was ∼13 times lower with Cu^2+^ added than without despite the abundance of *Methylococcaceae* family of methanotrophs (48.5% of 3.5±0.2×10^8^ eubacterial 16S rRNA copies mL^-1^).

## Discussion

Copper deficiency has been suggested as a potential cause for N_2_O emission from denitrification taking place in terrestrial and aquatic environments (33-35). In a previous study, Chang et al. demonstrated, using simplified pure-culture and co-culture experiments, that MB-OB3b lowered copper availability of the medium such that N_2_O reduction was substantially repressed in denitrifiers (30). Whether methanobactin-mediated N_2_O emission enhancement is relevant to actual soil environments with much more intricate microbiome complexity remained to be resolved, however. The results of the experiments in this study, although performed under well-controlled laboratory conditions, showed that methanobactin-enhanced N_2_O emissions may actually occur in soil environments with complex microbiomes. Methanobactin addition exerted significant influence on denitrification carried out by soils’ indigenous microbial consortia, demonstrating that diverse denitrifying community was still susceptible to methanobactin-mediated copper deprivation. Further, the N_2_O emission enhancement observed with the *Methylocystaceae* methanotroph-enriched reactor culture provided an unprecedented experimental evidence of indigenous alphaproteobacterial methanotrophs influencing N_2_O emission from denitrifiers. Observation of substantial *mbnA* transcription and the absence of this N_2_O emission enhancement effect in the copper-amended sample supported that methanobactin produced by the indigenous *Methylocystaceae* methanotrophs was likely the cause of the increased N_2_O production from denitrification. Additionally, the increased N_2_O production observed in soil enrichments with broadly varying *Methylocystaceae*-to-*Methlycoccaceae* ratios suggested that methanotrophic population composition *in situ* may be a major determinant of the copper-mediated N_2_O emission enhancement, while also suggesting existence of an additional mechanism via which methanotrophs affect N_2_O reduction in denitrifiers.

Evidence of the effect of methanobactin on the soil microbial consortia apart from N_2_O emission enhancement could also be discerned. Delays in NO_2_^-^ reduction during the progression of denitrification were observed in some, but not all, of denitrifying enrichments incubated in the presence of OB3b-MB capable of producing methanobactin. At least two distinct forms of nitrite reductases are known to mediate NO_2_^-^-to-NO reduction in denitrifiers: the copper-dependent nitrite reductase NirK and copper-independent cytochrome *cd*_1_ nitrite reductase NirS (36). In the previous investigation by Chang et al., indeed, NO_2_^-^ reduction by NirK-utilizing *Shewanella loihica*, but not NirS-utilizing denitrifiers, was affected by methanobactin-mediated Cu deprivation (30). The environmental conditions that select for enrichment of either NirK-or NirS-utilizing organisms remain unclear, and abundances of *nirS*-or *nirK*-possessing organisms vary in rice paddy soils (37). Possibly, the relative abundances and activities of the NirK- and NirS-utilizing denitrifiers may have varied in the denitrifying consortia prepared with soils A-E, explaining the non-uniform responses of the denitrifiers in the consortia to copper deprivation. That copper addition expedited NO_2_^-^ reduction in anoxic incubation of *nirK*-dominant chemostat culture was also in support of this hypothesis.

The shotgun metagenome analyses of the methanotrophic chemostat culture identified the organisms affiliated to the *Methylocystaceae* family as the dominant *nirK*-possessing organisms. Further, the *nosZ* gene affiliated to *Methylocystis* sp. SC2 was the dominant *nosZ* gene in the chemostat culture (38). These observations are certainly interesting, as the only *Methylocystaceae nirK* and *nosZ* sequences available in the database are the *nirK* gene found in the genome of *Methylocystis* sp. Rockwell and the *nosZ* gene found in a plasmid of *Methylocystis* sp. SC2 (38, 39). Both genes were unique, in a sense that they both shared <75% translated amino acid sequence identity with any other NirK or NosZ sequences in the NCBI database, and no previous study has reported recovery of these *Methylocystaceae nirK* and *nosZ* sequences in metagenome/metatranscriptome analyses. Despite the potential significance of the discovery, the possibility that the *Methylocystaceae* methanotrophs harboring these genes might have significantly affected NO_2_^-^ reduction and N_2_O production in the anoxic batch incubations was highly unlikely. Both *Methylocystis* sp. Rockwell and *Methylocystis* sp. SC2 have been confirmed of their inability to grow anaerobically, and neither strain has been confirmed of capability to reduce NO_2_^-^ or N_2_O utilizing NirK or NosZ, respectively (38, 39). Therefore, although the collected data are insufficient to completely rule out the possibility that these *Methylocystaceae* populations significantly contributed to denitrification and N_2_O production and consumption, it is more plausible that the N_2_O dynamics in the anoxic cultures were largely determined by non-methanotrophic facultatively anaerobic organisms carrying *nirK, nirS*, and/or *nosZ* genes.

In this metagenome analysis, many of the non-methanotrophic organisms putatively harboring *nirK* or *nirS* and those putatively harboring *nosZ* belonged to different taxa, suggesting that N_2_O production and N_2_O consumption were carried out by distinct organismal groups in the anoxic batch experiments performed with this chemostat culture. For example, the dominant non-*Methylocystaceae nirK* and *nosZ* recovered from the chemostat metagenome were those affiliated to *Rhizobiales* and *Bacteriodetes*, respectively, and only *nirK* and *nosZ* affiliated to *Pseudomonaceae* were recovered with similar coverage. These observations suggest that the alteration of copper availability brought about by methanobactin-producing methanotrophs may influence N_2_O emissions from denitrification occurring in modular manner, which may be the more likely case in complex environmental microbiomes (40, 41).

The distinctive contrast between the microbial compositions between the *mtri10* and *mtri10n* enrichments implied that methanobactin had pronounced impact on the overall microbial community. The microbial community formed from enrichment on CH_4_ and acetate after augmentation with the wildtype strain OB3b (*mtri10*) carried distinctively large proportion of *Moraxellaceae* (44.7%). Such abundance of *Moraxellaceae* was not observed in the enrichment with augmented Δ*mbnAN* mutant cells (*mtri10n*), suggesting that this enrichment of *Moraxellaceae* was due to the presence of methanobactin produced by *M. trichosporium* OB3b. The most probable explanations are selective bactericidal property and/or reduced copper bioavailability to competing organisms with high demands for copper (18, 22). The relative abundance of other phylogenetic groups including *Methylophilaceae* and *Flavobactericeae* also varied substantially across the treatments. Unlike the case for *Moraxellaceae*, however, the experimental evidences were not sufficient to attribute these alterations to the effect of methanobactin.

The observed N_2_O emission enhancement in methanotroph-enriched cultures dominated by the *Methylococcaceae* family, i.e., the *mtri0* enrichment, was unanticipated, as none of the sequenced genomes of the methanotrophs belonging to this phylogenetic group has been reported to produce methanobactin encoded by *mbnA* (18). What is evident from the experimental results, however, is that copper competition was central to N_2_O production in these enrichments, as copper amendment removed the N_2_O-accumulating phenotype. Indications that *Methylomicrobium album* BG8 and *Methylococcus capsulatus* Bath may utilize methanobactin-like copper chelators had been previously reported (42, 43). These *Methylococcaceae* strains tested positive on the Cu-CAS (chromo azurol S) plate assays, and the putative methanobactin-like compound of ∼1000 Da in size isolated from the spent medium of *M. capsulatus* Bath bound Cu with 1:1 stoichiometry; however, the genomic basis for synthesis of these compounds in *Methylococcaceae* methanotrophs has not yet been elucidated. Another copper uptake mechanism involving copper-binding periplasmic membrane proteins MopE and CorA have been identified in *M. capsulatus* Bath and *M. album* BG8, respectively (44, 45). The estimated binding constants of these putative copper chelators are tens of orders of magnitudes lower than those reported for methanobactin from *Methylocystaceae* methanotrophs; however, if present at large concentrations, the copper chelators may still exert significant impact on the Cu availability. Which, if any, of these mechanisms was responsible for copper withholding from denitrifiers and N_2_O reducers in the *Methylococcaceae*-dominant enrichments cannot be determined with the current data and warrants future investigation.

One of the most prominent characteristics of soil environments is their spatial and temporal heterogeneity, both in terms of physico-chemical makeup and microbial composition (46, 47). In CH_4_-enriched soil environments such as rice paddy and landfill soils, proteobacterial methanotrophs are often reported to be abundant, with *pmoA* copy numbers ranging between 10^7^ and 10^10^ gene copies g dry soil^-1^ (48-50). These numbers are of the same magnitude as the methanotrophic populations in the enrichment cultures observed to induce N_2_O production from the cohabiting denitrifying population in the laboratory experiments performed here. Methanotroph population density at local microsites may even be higher, especially at the oxic-anoxic interfaces, where CH_4_ and O_2_, the essential substrates of proteobacterial methanotrophs, are both available (49). Thus, it would not be surprising to find local concentrations of methanobactin-producing *Methylocystaceae* methanotroph communities in microniches within the soil sufficiently dense as to cause copper deficiency to co-habiting NosZ-utilizing N_2_O reducers. At microscopic scale, temporal oxic-to-anoxic shifts or *vice versa* would constantly occur at the oxic-anoxic interfaces, allowing for spatial co-existence of O_2_-dependent CH_4_ oxidation and O_2_-inhibited denitrification (32). The substrates of denitrification, NO_3_^-^ and NO_2_^-^, may be transported from oxic surface soils or supplied via oxidation of organic N or NH_4_^+^ *in situ* at the oxic-anoxic interface with intermittently available O_2_ (31). Periodic oxic-anoxic transitions may also take place in the vicinity of the water table in upland soils, as precipitation and drying cause fluctuations in the elevation of the water table (51). In such settings, aerobic microbial processes of nitrification and methanotrophy and anaerobic microbial processes of denitrification and methanogenesis may alternate at a larger scale. Snapshot-views of physico-chemical and biological states of soils may cast doubt on the likelihood of the methanotroph-enhanced N_2_O emissions occurring in actual soil environments; however, with these spatial and temporal shifts in consideration, the suggested mechanism is plausible in any terrestrial environment with high organic and nitrogen content. Thus, in approximating greenhouse gas budgets from environments such as rice paddy soils, landfill cover soils, wetland soils, and upland agricultural soils, N_2_O arising from the copper-mediated interaction between methanotrophs and N_2_O reducers need to be considered for development of a more accurate prediction model for N_2_O emissions.

## Methods and materials

### Soil sampling and characterization

Soil samples were collected from an experimental rice paddy located at Gyeonggido Agricultural Research & Extension Services in Hwaseong, Korea (37°13’21”N, 127°02’35”E) in August 2017 and four rice paddies near Daejeon, Korea (36°22’41”N, 127°19’50”E) in December 2018 (Supplementary Table S1). Samples were collected from the top layer of soil (0-20 cm below the overlying water). After removing plant material, soil samples were stored in sterilized jars, which were then filled to the brim with rice paddy water. The samples were immediately transported to the laboratory in a cooler filled with ice and stored at 4 °C until use. A small portion (∼50 g) of each collected soil was stored at −80 °C for DNA analyses.

The physicochemical characteristics of these soil samples were analyzed directly after sampling. The soil pH was measured by suspending 1 g wet weight soil in 5 mL Milli-Q water. Total nitrogen and carbon content of air-dried soil samples were analyzed with a Flash EA 1112 Elemental Analyzer (Thermo Fisher Scientific, Waltham, MA). For measurements of NH_4_^+^, NO_3_^−^, and NO_2_^−^ content in soil samples, 1 g air-dried soil was suspended in 5 mL 2 M KCl solution and shaken at 200 rpm for an hour. After settling the suspension for 10 min, the supernatant was passed through a 0.2 µM membrane filter (Advantec MFS Inc., Tokyo, Japan). The filtrate was analyzed using colorimetric quantification methods. The total copper content of soil samples was measured with an Agilent ICP-MS 7700S inductively coupled plasma mass spectrometer (Santa Clara, CA) after pretreatment with boiling aqua regia (52).

### Media and culture conditions

Unless otherwise mentioned, modified nitrate mineral salts (NMS) medium was used for enrichment of methanotrophs in soil microbial consortia and incubation of axenic cultures of the wildtype and Δ*mbnAN* mutant strains of *M. trichosporium* OB3b (53). The medium contained per liter, 1 g MgSO_4_·7H_2_O (4.06 mM), 0.5 g KNO_3_ (4.95 mM), 0.2 g CaCl_2_·2H_2_O (1.36 mM), 0.1 mL of 3.8% (w/v) Fe-EDTA solution (Sigma-Aldrich, St. Louis, MO), 0.5 mL of 0.02% (w/v) Na_2_MoO_4_·2H_2_O solution, and 0.1 mL of the 10,000X trace element stock solution (Supplementary Table S5). All glassware used in this study was equilibrated in a 5 N HNO_3_ acid bath overnight before use to reduce background contamination of copper to <0.01 µM in media. Forty-milliliter aliquots of the medium was distributed into 250-mL serum bottles and the bottles were sealed with butyl-rubber stoppers (Geo-Microbial Technologies, Inc., Ochelata, OK, US) and aluminum caps. After autoclaving, the media were amended with 0.2 mL of 200X Wolin vitamin stock solution (Supplementary Table S6) and 500 mM pH 7.0 KH_2_PO_4_/Na_2_HPO_4_ solution was added to a final concentration of 5 mM (54). High-purity CH_4_ (>99.95 %; Deokyang Co., Ulsan, Korea) replaced 20% of the headspace air. After inoculation, culture bottles were incubated in dark at 30 °C with shaking at 140 rpm.

### Analytical procedures

Headspace CH_4_ concentrations were measured using a GCMS-QP2020 gas chromatograph-mass spectrometer (Shimadzu Cooperation, Kyoto, Japan) equipped with a SH-Rt™-Q-BOND column (30 m length × 0.32 mm inner diameter, 10 μm film thickness). The injector and oven temperatures were set to 150 °C and 100 °C, respectively, and helium was used as the carrier gas. Headspace N_2_O concentrations were measured with a HP 6890 Series gas chromatograph equipped with a HP-PLOT/Q column and an electron capture detector (Agilent, Palo Alto, CA, US). The injector, oven, and detector temperatures were set to 200, 85, and 250 °C, respectively. For each CH_4_ or N_2_O measurement, 50 μL or 100 μL of gas sampled using a Hamilton 1700-series gas-tight syringe (Reno, NV) was manually injected into the gas chromatographs. The dissolved N_2_O concentrations were calculated from the headspace concentrations using the dimensionless Henry’s law constant of 1.92 at 30°C (55, 56). The dissolved concentrations of NO_3_^-^ and NO_2_^-^ were determined colorimetrically using the Griess reaction (57). As the assay measures NO_2_^-^, vanadium chloride (VCl_3_) was used to reduce NO_3_^-^ to NO_2_^-^. The concentration of NH_4_^+^ was measured using the salicylate method (58).

The total bacterial population was quantified using TaqMan-based quantitative polymerase chain reactions (qPCR) targeting eubacterial 16S rRNA gene (1055f: 5’-ATGGCTGTCGTCAGCT-3’; 1392r: 5’-ACGGGCGGTGTGTAC-3’; and Bac1115_probe: 5’-CAACGAGCGCAACCC-3’)(59). The primers and probe set exclusively targeting the *pmoA* gene (encoding for the β subunit of particulate methane monooxygenase) of *M. trichosporium* OB3b (OB3b_pmoAf: 5’-CGCTCGACATGCGGATAT-3’; OB3b_pmoAr: 5’-TTTCCCGATCAGCCTGGT-3’; and OB3b_ pmoA_probe: 5’-AGCCACAGCGCCGGAACCA-3’) was designed *de novo* using Geneious v9.1.7 software (Biomatters Ltd., Auckland, New Zealand) and used for quantification of *M. trichosporium* OB3b cells. qPCR assays were performed with a QuantStudio™ 3 Real-Time PCR System (Thermo Fisher Scientific). The calibration curves for the targeted genes were constructed using serial dilutions of PCR2.1™ vectors (Thermo Fisher Scientific) carrying the target fragments. For each qPCR quantification, three biological replicates were processed separately. The complete list of the primers and probes used in this study are provided in Table 1.

**Table 1.**
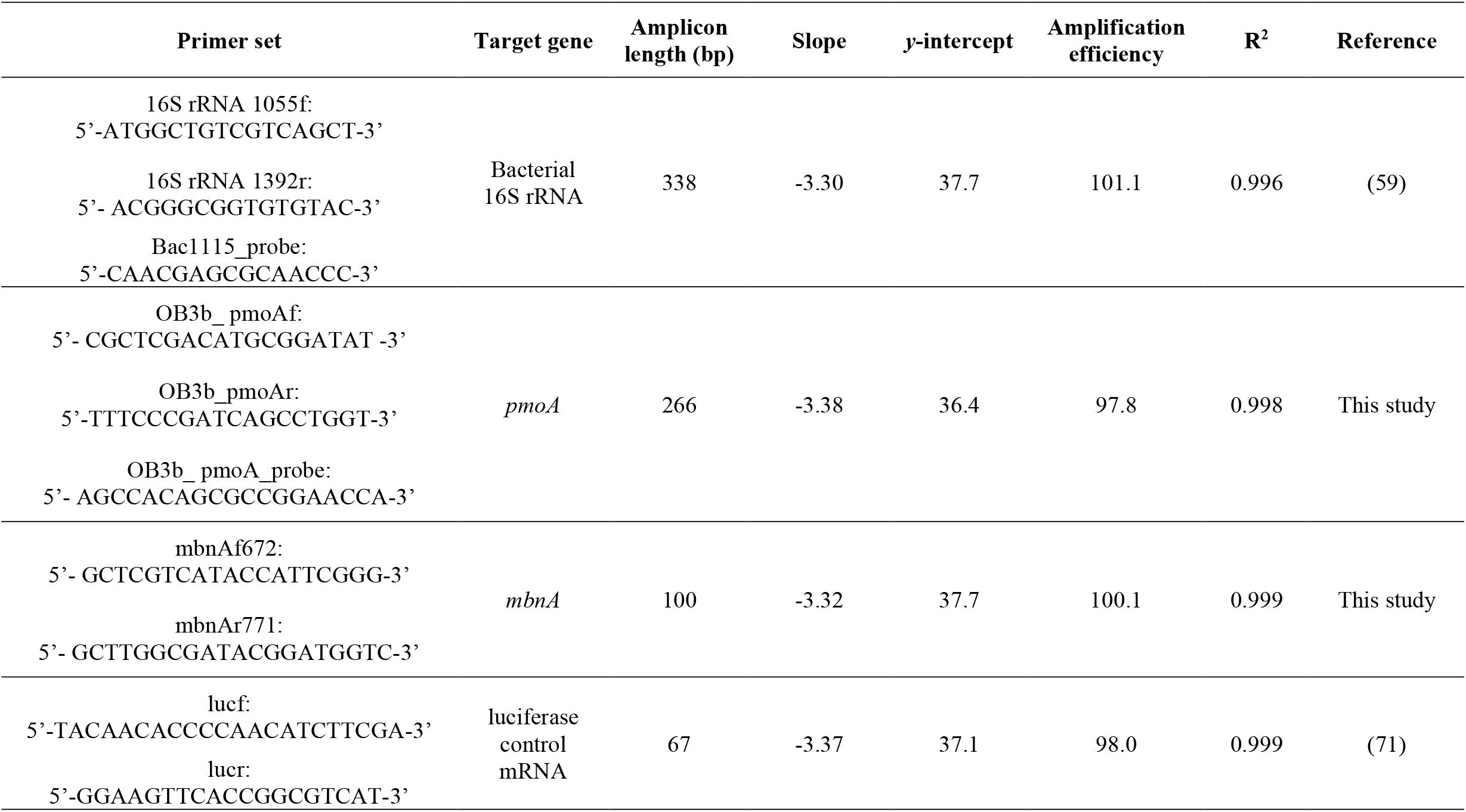
List of the primers/probe sets used for qPCR and RT-qPCR analyses

### Monitoring of denitrification and N_2_O production in rice paddy soil suspensions amended with methanobactin from *M. trichosporium* OB3b

Methanobactin was isolated from *Methylosinus trichosporium* OB3b cultures and purified as previously described (60). The effects of MB-OB3b on N_2_O emissions were examined with suspensions of five rice-paddy soils (soil A-E) with varying copper content. Each soil suspension was prepared by suspending 1 g of air-dried rice paddy soil into 50 mL Milli-Q water in 160-mL serum bottles and the headspace was replaced with N_2_ gas after sealing. Methanobactin stock solution was prepared by dissolving 10 µmoles (11.5 mg) freeze-dried methanobactin in 10 mL Milli-Q water. The stock solution was equilibrated in an anaerobic chamber filled with 95% N_2_ and 5% H_2_ (Coy Laboratory Products Inc., Grass Lake, MI) for one hour to remove dissolved O_2_, and 0.5 mL of the solution was added to the soil suspensions. Potassium nitrate and sodium acetate were then added to final concentrations of 5 mM and 10 mM, respectively. Soil suspensions were incubated with shaking at 140 rpm at 30°C. The headspace N_2_O concentration and the dissolved concentrations of NO_3_^-^ and NO_2_^-^ were monitored until depletion of NO_3_^-^ and NO_2_^-^. The dissolved concentrations of NH_4_^+^ were also monitored to ensure absence of significant dissimilatory nitrate reduction to ammonium (DNRA) activity. Negative controls were prepared identically, but without methanobactin amendment. Abiotic control experiments with sterilized soils with and without methanobactin amendment were performed with 5 mM NO_3_^-^ or NO_2_^-^ added, to confirm that abiotic contribution to N_2_O production was minimal.

### Monitoring of denitrification and N_2_O production in subcultures extracted from a quasi-steady state chemostat culture of *Methylocystaceae*-dominated soil methanotrophs

A continuous methanotrophic enrichment culture was prepared with Soil E. The inoculum was generated by suspending 1 g (wet wt) soil in 50 mL NMS medium in a sealed 160-mL serum vial with air headspace. The vials were fed twice with 41 μmoles CH_4_, yielding a headspace concentration of ∼0.5% v/v upon each injection. After the initial batch cultivation, 20 mL of the enrichment was transferred to 5 L NMS medium in a 6-L fermenter controlled with a BioFlo^®^ 120 Bioprocess Control Station (Eppendorf, Hamburg, Germany). Gas stream carrying a relatively low CH_4_ concentration (0.5% v/v in air) was fed continuously at a rate of 145 mL min^-1^ through a gas diffuser to stimulate growth of *Methylocystaceae* methanotrophs, as previous fed-batch and chemostat incubation of rice paddy soils with a continuous supply of 20% v/v CH_4_ resulted in domination by gammaproteobacterial methanotrophs (61). The reactor culture was maintained in the dark at pH 6.8 and 30 °C with agitation at 500 rpm. After the initial fed-batch operation, the reactor culture was transitioned to continuous operation with the dilution rate set to 0.014 h^-1^. After quasi-steady state was attained, as indicated by constant effluent cell density and CH_4_ concentration, 1.0-mL aliquots were collected and the pellets were stored at −20 °C for analyses of the microbial community composition and denitrification functional genes and qPCR quantification of *mbnA* and 16S rRNA genes. Additionally, 0.4-mL aliquots were treated with RNAprotect Bacteria Reagent (Qiagen, Hilden, Germany) and stored at −80 °C for reverse transcription qPCR (RT-qPCR) analyses of *mbnA* expression.

Triplicate anoxic batch cultures were prepared by distributing 50 mL culture harvested from the running quasi-steady state reactor to 160-mL serum bottles and flushing the headspace with N_2_ gas. A denitrifying enrichment was prepared separately by incubating 1 g (wet wt) of Soil E in 100-mL anoxic NMS medium amended with 10 mM sodium acetate. One milliliter of this denitrifier inoculum was added to the methanotroph enrichments, to which 20 mM sodium acetate was added as the electron donor for denitrification. The cultures were amended with or without 2 μM CuCl_2_ and the culture vessels were incubated in the dark with shaking at 140 rpm at 30 °C and the changes to the amounts of NO_3_^-^, NO_2_^-^, and N_2_O-N in the vessels were monitored until no further change was observed.

### Denitrification and N_2_O production in soil enrichments with altered methanotrophic population composition

Varying amounts of pre-incubated *M. trichosporium* OB3b cells were added to rice paddy soil slurry microcosms before enrichment with CH_4_, to artificially vary the proportion of the methanobactin-producing subgroup among the total methanotroph population. The total indigenous gammaproteobacterial methanotroph population in the soil suspension was approximated by multiplying the total eubacterial 16S rRNA copy number per mL suspension (as determined with qPCR) by the proportion of the 16S rRNA gene sequences affiliated to gammaproteobacterial methanotrophs (all affiliated to the *Methylococcaceae* family) in soil “B” (as computed from the MiSeq amplicon sequencing data, Supplementary Table S7). The number of *M. trichosporium* OB3b cells in the precultures were estimated with the TaqMan qPCR targeting the duplicate *pmoA* genes of the strain. The *M. trichosporium* OB3b preculture was used to prepare soil suspensions with augmented *Methylocystaceae* population. The ratios of *M. trichosporium* OB3b *pmoA* copy numbers to the estimated 16S rRNA gene copy numbers of the *Methylococcaceae* methanotrophs were set to 1 and 10 in the 40 mL suspensions. It should be stressed that these ratios cannot be directly translated to cell number ratios, as the numbers of 16S rRNA genes in the completed genomes of gammaproteobacterial methanotrophs deposited in NCBI GenBank database vary between one and four. Soil suspensions without *M. trichosporium* OB3b augmentation was also prepared. As a negative control, the Δ*mbnAN* mutant strain of *M. trichosporium* OB3b was added in place of the wildtype strain OB3b (62). A copper-replete control was also prepared for the experimental condition with no *M. trichosporium* OB3b augmentation by amending a subset of cultures with 2 μM CuCl_2_.

The soil suspensions were prepared by adding 0.1 g wet wt of the soil to the autoclaved modified NMS medium bottles prepared as described above (40 mL NMS medium in 250-mL serum bottles). After sealing the bottles and adding *M. trichosporium* OB3b or Δ*mbnAN* mutant preculture or CuCl_2_ to the calculated target concentrations, 20% of the headspace was replaced with CH_4_ and sodium acetate was added to a final concentration of 5.0 mM. When the headspace CH_4_ concentration decreased to 10%, the headspace was replenished with an 80:20 mixture of air and CH_4_, and the cultures were amended with 200 μmoles NO_3_^-^ and 400 μmoles sodium acetate. At each sampling time point, the headspace N_2_O concentration was measured and 0.5 mL of the culture collected from each bottle for determination of NO_3_^-^, NO_2_^-^, and NH_4_^+^ concentrations. The cell pellets collected immediately after CH_4_ concentration dropped to ∼10% were subjected to qPCR targeting strain OB3b *pmoA* genes and eubacterial 16S rRNA genes. The 16S rRNA amplicon sequences of the pellets were also analyzed.

### Microbial community composition analyses and prediction of genomic potentials for denitrification reactions from 16S rRNA amplicon sequences

Genomic DNA was extracted with DNeasy PowerSoil Kit (Qiagen, Hilden, Germany). The V6-V8 region of the 16S rRNA gene amplified with the universal primers 926F (5’-AAACTYAAAKGAATTGRCGG - 3′)/1392R (5’-ACGGGCGGTGTGTRC-3′) was sequenced at Macrogen Inc. (Seoul, Korea) using a MiSeq sequencing platform (Illumina, San Diego, CA)(63). The raw sequence data were processed using the QIIME pipeline v1.9.1 with the parameters set to default values (64). After demultiplexing and quality trimming, the filtered sequences were clustered into operational taxonomic units (OTUs) using USEARCH algorithm, and the OTUs were assigned taxonomic classification using the RDP classifier against the Silva SSU database release 132. The raw sequences were deposited to the NCBI SRA database (SRX8210313, SRX8209287-8209292).

### Metagenomic analysis of the chemostat enrichment and reverse-transcription (RT) qPCR targeting *mbnA*

Shotgun metagenomic sequencing of the reactor culture was performed at Macrogen Inc. using a HiSeq X sequencing platform (Illumina) generating ∼5 Gb of paired-end reads with 150-bp read length (accession number: SRX8210314). The raw short reads were quality-screened using Trimmomatic v0.36 with default parameters (65). The processed reads were assembled into contigs using metaSPAdes v3.12.0, and coding sequences were identified using Prodigal v2.6.3 (66, 67). The Hidden Markov model (HMM) algorithms for *nirS, nirK* and clade I and clade II *nosZ* were downloaded from the FunGene database (http://fungene.cme.msu.edu/). A previously constructed HMM algorithm for *mbnA* was used for screening of *mbnA* genes (68). The predicted contigs were screened for these targeted genes using *hmmsearch* in HMMER 3.1b2 with E-value set to 10^−5^ using these HMM algorithms. The identified gene sequences were further validated by running blastx against the RefSeq database with E-value cut-off set to 10^−3^. For taxonomy assignment, the translated amino acid sequences of the screened functional genes were searched against NCBI’s non-redundant protein database (nr) using blastp with E-value cutoff of 10^−10^. The quality-trimmed reads were mapped onto the screened functional gene sequences using the bowtie2 v2.3.4.1 software with the parameters set to default values (69). The coverage of each target gene was calculated using the *genomecov* command of the BEDtools v2.17.0 software and normalized by its length (70).

A degenerate primer set was designed with the sole unique *mbnA* sequence recovered from the shotgun metagenome data and publicly available *Methylocystis mbnA* sequences with >85% translated amino acid sequence identity with this metagenome-derived *mbnA* sequence (Supplementary Fig. S1) using Geneious v9.1.7 software (Biomatters Ltd., Auckland, New Zealand). This primer set (672f: 5’-GCTCGTCATACCATTCGGG-3’ and 771r: 5’-GCTTGGCGATACGGATGGTC-3’) was used for quantification of *mbnA* genes and transcripts in the chemostat culture (Supplementary Fig. S1). Extraction and purification of RNA was performed using RNeasy Mini Kit (Qiagen) and RNA MinElute Kit (Qiagen), respectively, and reverse transcription was performed using SuperScript III reverse transcriptase (Invitrogen, Carlsbad, CA), according to the established protocols (71). As previously described, a known quantity of luciferase control mRNA (Promega, Madison, WI) was used as the internal standard to correct for RNA loss during extraction and purification procedure. The *mbnA* transcript copy numbers were normalized with *mbnA* copy numbers in the genomic DNA.

### Statistical analyses

All experiments were performed in triplicates and the average values were presented with the error bars representing the standard deviations of triplicate samples. The statistical significance of the data from two different experimental conditions were tested with the two-sample *t*-tests, and statistical comparison between two different time points with one-sample *t*-tests. The SPSS statistics 24 software (IBM Corp, NY) was used for statistical analyses.

## Acknowledgements

This work was supported by the National Research Foundation of Korea (Grants 2017R1D1A1B03028161 and 2015M3D3A1A01064881) and the United States Department of Energy (DE-SC0020174). The authors were also financially supported by the Brain Korea 21 Plus Project (Grant 21A20132000003).

